# Somatic mutations reveal widespread mosaicism and mutagenesis in human placentas

**DOI:** 10.1101/2021.01.26.428217

**Authors:** Tim H. H. Coorens, Thomas R. W. Oliver, Rashesh Sanghvi, Ulla Sovio, Emma Cook, Roser Vento-Tormo, Muzlifah Haniffa, Matthew D. Young, Raheleh Rahbari, Neil Sebire, Peter J. Campbell, D. Stephen Charnock-Jones, Gordon C. S. Smith, Sam Behjati

## Abstract

Clinical investigations of human fetuses have revealed that placentas occasionally harbour chromosomal aberrations that are absent from the fetus^1^. The basis of this genetic segregation of the placenta, termed confined placental mosaicism, remains unknown. Here, we investigated the phylogeny of human placentas reconstructed from somatic mutations, using whole genome sequencing of 86 placental biopsies and of 106 microdissections. We found that every placental biopsy represented a clonal expansion that is genetically distinct. Biopsies exhibited a genomic landscape akin to childhood cancer, in terms of mutation burden and mutational imprints. Furthermore, unlike any other human normal tissue studied to date, placental genomes commonly harboured copy number changes. Reconstructing phylogenetic relationships between tissues from the same pregnancy, revealed that developmental bottlenecks confined placental tissues, by separating trophectodermal from inner cell mass-derived lineages. Of particular note were cases in which inner cell mass-derived and placental lineages fully segregated within a few cell divisions of the zygote. Such early embryonic bottlenecks may enable the normalisation of zygotic aneuploidy. We observed direct evidence for this in a case of mosaic trisomic rescue. Our findings reveal cancer-like mutagenesis in placental tissues and portray confined mosaicism as the normal outcome of placental development.

## INTRODUCTION

The human placenta is a temporary organ whose dysfunction contributes substantially to the global burden of disease^2^. Amongst its many peculiarities is the occurrence of chromosomal aberrations confined to the placenta, which are absent from the newborn infant. First described by Kalousek and Dill in 1983^1^, confined placental mosaicism affects one to two percent of pregnancies^3^. It may pervade both components of placental villi, the trophectoderm or the inner cell mass-derived mesenchyme, alone or in combination.

The genetic segregation of placental biopsies in confined placental mosaicism suggests that bottlenecks may exist in early development that provide opportunities for genetically separating placental and fetal lineages. It is conceivable that these are physiological genetic bottlenecks underlying the normal somatic development of placental tissue. Alternatively, genetic segregation may represent pathological perturbation of the normal clonal dynamics of early embryonic lineages. For example, it has been suggested that confined placental mosaicism represents a depletion from the fetus-forming inner cell mass of cytogenetically abnormal cells, commonly found in early embryos^4^.

The clonal dynamics of human embryos cannot be studied prospectively. It is, however, possible to reconstruct embryonic lineage relations from somatic mutations that had been acquired during cell divisions, serving as a record of early embryonic lineage relations^5–8^. Furthermore, these mutations may reveal specific mutagenic processes that shape a tissue^9^. Here, we studied the somatic genetic architecture of human placentas by whole genome sequencing, to investigate the clonal dynamics and mutational processes underpinning the development of human placentas.

## RESULTS

### Somatic mutations in placental biopsies

The starting point of our investigation were whole genome sequences of 86 placental biopsies, obtained from 37 term placentas along with inner cell mass derived umbilical cord tissue and maternal blood (**Fig. 1a**). Tissues had been curated by the Pregnancy Outcome Prediction study, a prospective collection of placental tissue and extensive clinical data, including histological assessment of individual biopsies, described in detail elsewhere^10^. We included placentas from normal pregnancies and from pregnancies associated with a range of abnormal parameters (Extended Data Table 1). Placental and umbilical cord biopsies were washed in phosphate-buffered saline to remove maternal blood. We assessed the possibility of residual contamination of biopsies with maternal blood by searching biopsy DNA sequences for germline polymorphisms unique to the mother (Extended Data Fig.1).

**Figure 1.**
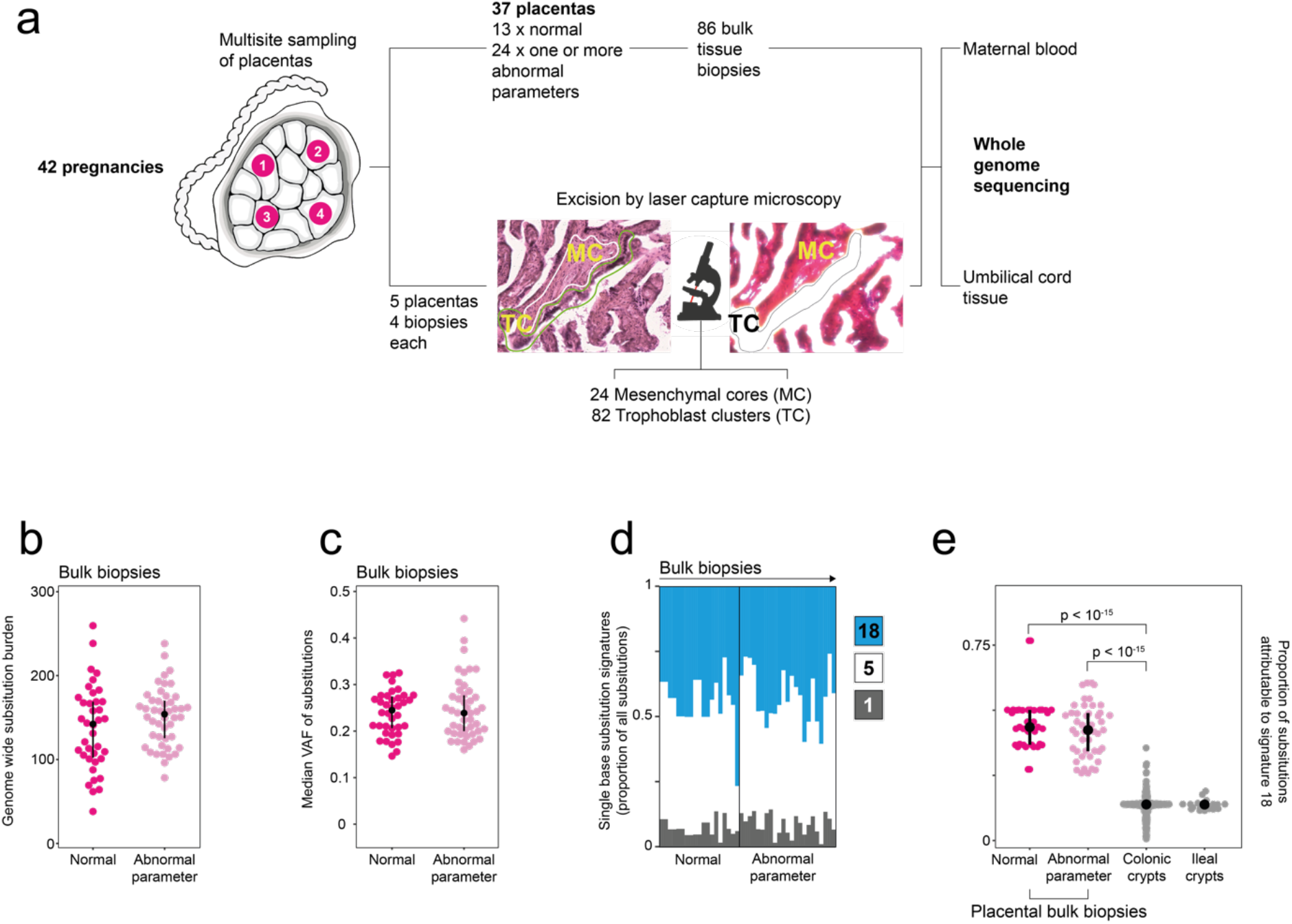
The genomes of placental bulk biopsies. (**a**) Workflow detailing experimental design with photomicrograph demonstrating microdissection of trophoblast. (**b**) Substitution burden per placental biopsy, adjusted for coverage and median VAF. An abnormal pregnancy is defined by the deviation of one or more clinically validated markers from their normal range over the course of pregnancy (Extended Data Table 1). (**c**) Median variant allele frequency of substitutions in each placental biopsy. (**d**) Single base substitution signatures in placental biopsies. Each column represents one biopsy. Colours represent signatures, as per legend. (**e**) Prevalence of signature 18 mutations in placental bulk biopsies in comparison to human intestinal tissue^12^, the normal tissue with the highest prevalence of signature 18 variants reported to date.

We called substitutions in placental biopsies and found a high burden of mutations (**Fig. 1b**). This was an unexpected result because we had assumed that the placenta – a normal, non-cancerous bulk tissue – was polyclonal. Examining polyclonal tissues by whole genome sequencing does not usually reveal somatic mutations, with the exception of a small number of non-heterozygous post-zygotic (mosaic) variants that represent cell divisions of the early embryo^5–8^. However, in placental biopsies we found a mean of 145 base substitutions per biopsy (range 38-259). The average median variant allele frequency (VAF) of placental mutations within each biopsy was 0.24 (range 0.15-0.44) which indicated the mutations pervaded on average ~50% of cells (**Fig. 1c**).

Base substitutions can be classified by their trinucleotide context into mutational signatures, which may reveal mutagenic processes that shaped a tissue^9^. Accordingly, three different single base substitution mutational signatures characterized substitutions of placental biopsies: signatures 1, 5, and 18 (**Fig. 1d**). Signatures 1 and 5 are ubiquitous in human tissues and accumulate throughout life^9^. In contrast, signature 18 variants, which may be associated with reactive oxygen species and oxidative stress^11^, are seen infrequently in normal tissues. In placental biopsies, signature 18 contributed ~43% of substitutions. In comparison, in normal human colorectal crypts, the normal tissue with the highest prevalence of signature 18 mutations described to date^12^, it contributed an average of ~13% of substitutions (**Fig. 1e**).

Other classes of somatic mutations, small insertions and deletions (indels) and copy number changes (Extended Data Table 2), confirmed the clonal composition of biopsies. Of note, 41/86 biopsies harboured at least one copy number change (gain or loss; median size per unique segment, 73.6 kb). However, only one aberration, a trisomy of chromosome 10, would have been detectable by clinical karyotyping of chorionic villi. Within the constraints of the sample size of each clinical group, we did not observe systematic differences in overall mutation burden and spectra between groups (Extended Data Fig.2). Comparing somatic changes between multiple biopsies from the same placenta showed that the majority were unique to the given sample, suggesting that each biopsy represented a genetically independent unit (Extended Data Fig.3). Of note, placental biopsies had been obtained from separate quadrants of the placenta, several centimetres apart, thus representing distinct lobules. These observations indicated, therefore, that placental biopsies inherently possessed confined, mosaic genetic alterations.

### Monoclonal organization of trophoblast clusters underpins mosaicism of biopsies

To investigate the cellular origin of the mosaicism we observed at the level of placental biopsies, we directly assessed the genomes of the two main elements comprising chorionic villi, namely inner-cell mass derived fetal mesenchymal cores and the trophoblast (**Fig. 1a**). Using laser capture microscopy we excised 82 trophoblast clusters and 24 mesenchymal cores from the term placentas of five normal pregnancies and subjected these to whole genome sequencing. We first called substitutions unique to each trophoblast cluster or mesenchymal core, and assessed their VAF distribution. If groups of cells were organized as monoclonal patches derived from a single stem cell, their mutations would exhibit a VAF close to 0.5, as for example has been observed in single colonic crypts or single endometrial glands (**Fig. 2a**). Alternatively, if groups of cells were of oligo- or polyclonal origin, their median VAF would be shifted towards zero (**Fig. 2a**). We found the median VAF of trophoblast clusters and mesenchymal cores significantly differed (0.39 versus 0.20, Wilcoxon rank sum test, p < 10^−^ 12) (**Fig. 2a**). This indicated that trophoblast clusters exhibited a VAF distribution consistent with a monoclonal architecture, whereas mesenchymal cores did not. Hence, the mosaicism observed in bulk biopsies emanated from the trophoblast.

**Figure 2.**
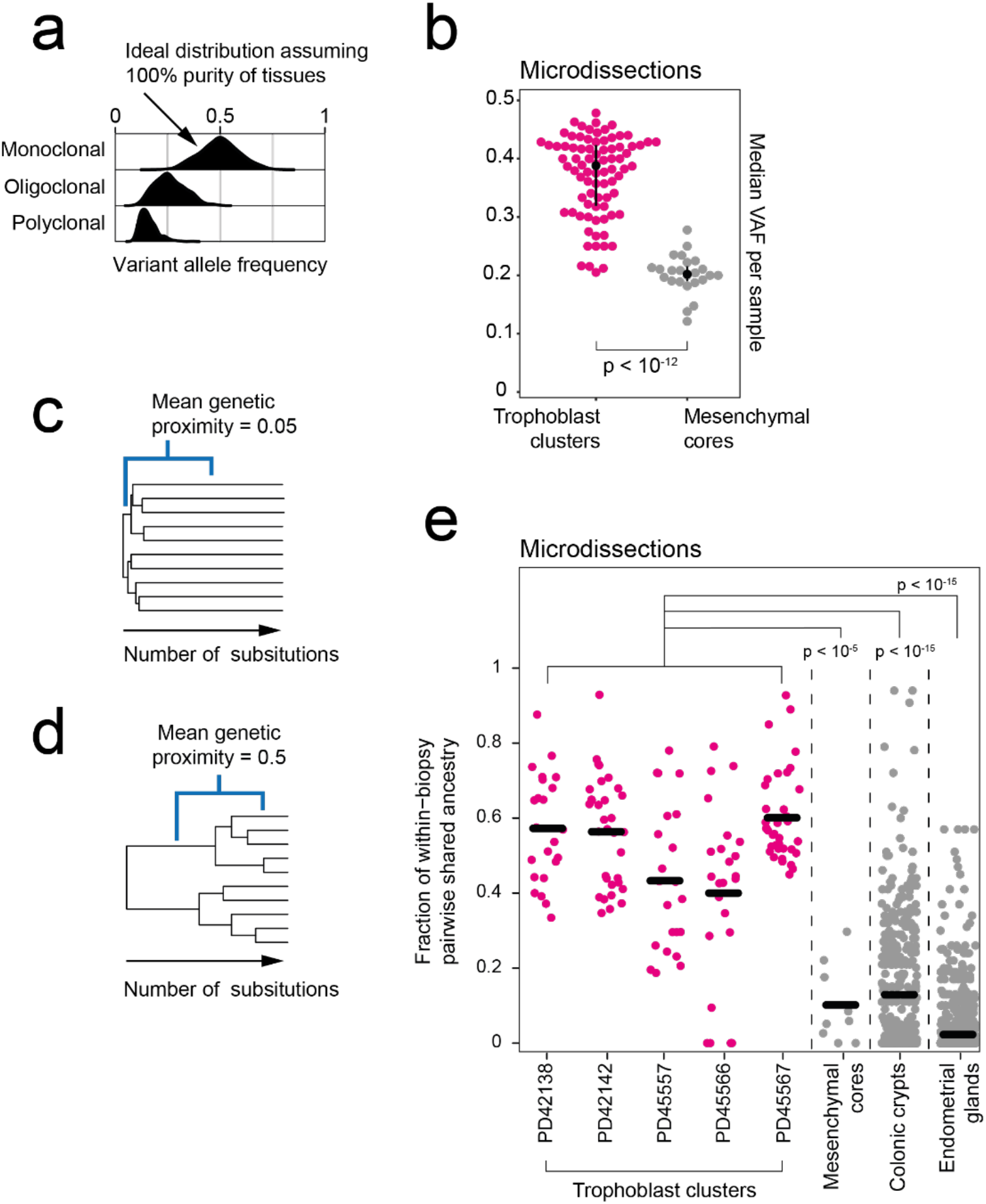
Clonal architecture of microdissected trophoblast clusters and mesenchymal cores. (**a**) Theoretical, expected VAF distribution as per different clonal architecture, assuming 100% purity. (**b**) Comparison of the median substitution VAF between microdissected trophoblast and mesenchymal cores. P-value refers to the Wilcoxon rank sum test comparing the two groups. (**c,d**) Genetic proximity scores were calculated as the fraction of shared mutations of a pair of samples divided by their mean total mutation burden. For example, a mean score of 0.05 conveys little sharing (**c**), while 0.5 signifies a longer shared development (**d**). (**e**). Genetic proximities across trophoblast clusters and mesenchymal cores from the same placental biopsies and data from colonic crypts^12^ and endometrial glands^13^. Each dot represents the comparison of two of the same histological unit (e.g., two trophoblast clusters) from the same biopsy. To avoid including adult clonal expansions, bifurcations in phylogenies after 100 post-zygotic mutations were not considered for colon and endometrium. P-values refer to assessment by Wilcoxon rank sum test.

We further corroborated this conclusion by studying the genetic relationship between trophoblast derivatives and mesenchymal cores from the same biopsies. We constructed phylogenetic trees and calculated pairwise genetic proximity scores of microdissections of the two components. We defined this score as the fraction of shared mutations out of the total mutation burden of the pair. A low genetic proximity score for pairs of trophoblast clusters or of mesenchymal cores from the same biopsy would indicate that the pool of precursor cells forming these diverged early in development (**Fig. 2c**). By contrast, a high score would suggest that histological units within each patch of tissue arose from only a few precursor cells with a relatively long shared ancestry (**Fig. 2d**). This analysis revealed a significant difference in the developmental clonal composition between trophoblast clusters and mesenchymal cores (p < 10^−5^; Wilcoxon rank sum test) (**Fig. 2e**). On average, within each biopsy, pairs of trophoblast clusters shared 53% of somatic mutations, indicating a long, joint developmental path of these cells. In contrast, pairs of mesenchymal cores from the same biopsy exhibited a mean genetic proximity of 10% and thus a short, shared phylogeny, in line with other inner cell mass-derived tissues, such as colon and endometrium^12,13^ (**Fig. 2e**). These observations suggest that large expansions of single trophoblastic progenitors underpin the normal clonality and confined mosaicism of placental biopsies we observed.

### Biases in cell allocation to trophectoderm and inner cell mass

Our findings thus far indicated that the seeding of a patch of placental tissue represented a genetic bottleneck at which clinically detectable, trophoblastic confined placental mosaicism could arise. We now considered whether earlier bottlenecks may exist prior to seeding of the placenta, amongst the first cell divisions of the embryo. Accordingly, we assessed the distribution of early embryonic lineages across placental and inner cell mass derived tissues by measuring the VAFs of post-zygotic (early embryonic) mutations, representing the first cell divisions of the zygote (**Fig. 3a and Fig. 3b**). These are mutations present in umbilical cord or placenta which, unlike heterozygous germline variants, present at a variable VAF across tissues.

**Figure 3.**
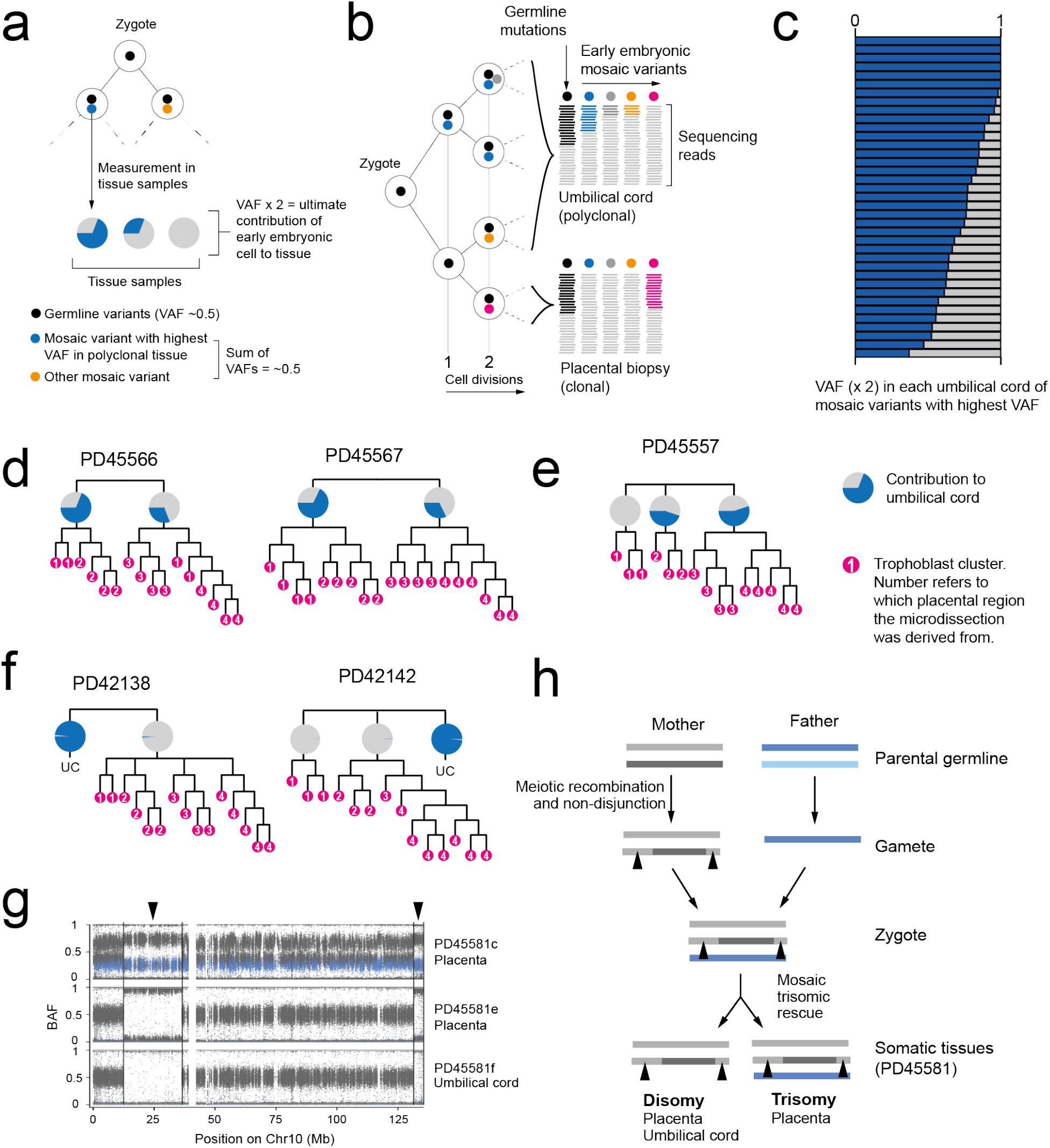
Early embryonic genetic bottlenecks and their relationship to trisomic rescue. ( a) Schematic depicting the detection of the earliest post-zygotic mutations and the estimation of contribution to samples from their variant allele frequencies. (**b**) Hypothetical lineage tree of early embryo showing how measurements of VAF may relate to cell divisions. (**c**) The contribution of the major lineage to the umbilical cord as calculated from the embryonic mutation with the highest VAF. (**d**) Early trees of trophoblast clusters of PD45566 and PD45567, with the contribution of lineages to the umbilical cord coloured in blue in pie charts. The umbilical cord exhibits an asymmetric contribution of the daughter cells of the zygote. (**e**) Early cellular contribution in PD45557 shows separation of one placental lineage. (**f**) In PD42138 and PD42142 the placental and umbilical cord lineages do not share any early embryonic mutations. (**g**) B-allele frequency (BAF) of germline SNPs on chromosome 10 in PD45581, showing a trisomy in PD45581c. SNPs absent from mother are coloured in blue. (**h**) Overview of genomic events in PD45581 and parents leading to the observed mosaic trisomic rescue. The arrowheads highlight areas of two genotypes in PD45581c due to meiotic recombination in the mother.

We directly compared the VAF of early embryonic mutations across biopsies and microdissected tissues, examining a total of 234 samples from 42 pregnancies. We found three configurations that identified two early embryonic bottlenecks (**Fig. 3, c to f**). In about half of pregnancies (19/42), the earliest post-zygotic mutation exhibited an asymmetric VAF across inner cell mass and trophectoderm lineages, without genetically segregated placental samples in this configuration (**Fig. 3d,** Extended Data Fig.4). In about a quarter of pregnancies (12/42), we found that one placental biopsy did not harbor the early embryonic mutations shared between umbilical cord and other placental biopsies. This indicated that the primordial cell seeding the placental biopsy in question segregated in early embryogenesis, thus representing a genetic bottleneck (**Fig. 3e,** Extended Data Fig.5). Loss of heterozygosity as an explanation for the absence of early embryonic mutations was excluded (Extended Data Table 4). In the remaining quarter (11/42) of pregnancies, the genetic bottleneck generated a complete separation of all placental tissues from umbilical cord samples **(Fig. 3f,** Extended Data Fig.6). There were no shared mutations, including early embryonic mutations, between placental tissues and umbilical cord lineages, consistent with this complete split having occurred at the first cell division of the zygote. Taken together, this data suggests that in about half of placentas, at least one bottleneck exists. Consequently, genomic alterations that pre-exist in the zygote, or arise within the first few cell divisions, may segregate between placenta and fetal lineages.

### Trisomic rescue through an early embryonic genetic bottleneck

A striking example of segregating genomic alterations that pre-exist in the zygote was a pregnancy harboring trisomy of chromosome 10 in one placental biopsy, but disomy of chromosome 10 elsewhere in the placenta and umbilical cord (**Fig. 3g**). Analysis of the distribution of parental alleles demonstrated that there were two maternal and one paternal chromosome in the affected placental biopsy. Importantly, the two maternal copies were non-identical, generating segments of chromosome 10 with three genotypes in the affected placental biopsy. In samples which were disomic for chromosome 10, there were two maternal copies, i.e. uniparental (maternal) disomy (**Fig. 3h**). This pattern demonstrates direct evidence for trisomic rescue, i.e. that the trisomy was present in the zygote, but that one cell of the two-cell embryo, which ultimately formed the fetus and some of the placenta, reverted to disomy post-zygotically (**Fig. 3h**). As the extra chromosome was maternal and the chromosome lost paternal, the fetus was euploid with uniparental (maternal) disomy. The pattern of the VAF distribution of early embryonic mutations across all tissues obtained from this pregnancy indicated that the trisomic rescue had occurred at a genetic bottleneck within the first cell divisions (Extended Data Fig.6).

### Mutational landscape of trophoblast clusters

The monoclonal organization of trophoblast clusters provided the opportunity to examine mutational processes that forged placental tissue in detail. Examining base substitutions of individual trophoblast clusters further, we found an average of 192 variants per cluster (Extended Data Fig.7). The mutation rate of trophoblast clusters was similar to that of childhood cancers, which, like the placenta, are primarily shaped by the mutational processes of fetal life^14^ (**Fig. 4a**). Furthermore, a large proportion of substitutions in each trophoblast sample could be assigned to signature 18 (**Fig. 4b and Fig. 4c**), exceeding what has been observed in rhabdomyosarcoma and neuroblastoma, the cancer types with the highest relative burden of signature 18 variants^14^ (**Figure 4c**). In addition, we found an indel burden proportional to substitutions in each sample, as well as widespread copy number changes (Extended Data Fig.8, Extended Data Tables 2 and 4).

**Figure 4.**
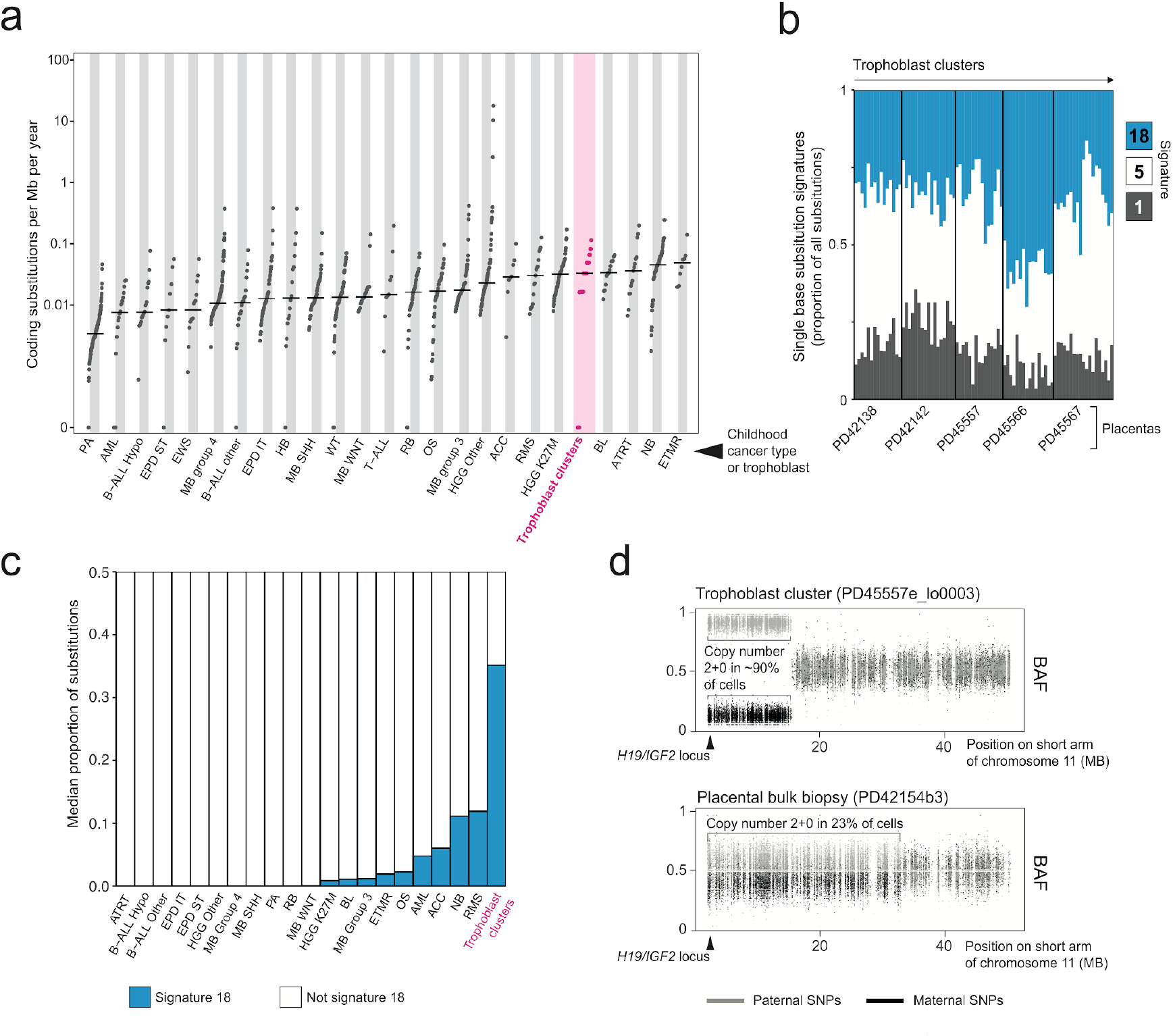
The genomes of microdissected trophoblast clusters. (**a**) Comparison of the coding substitutions per Mb per year between trophoblast microdissections and a range of paediatric malignancies^14^. Per year estimates are corrected for gestation. Abbreviations: Pilocytic astrocytoma (PA), acute myeloid leukaemia (AML), hypodiploid B-cell acute lymphoblastic leukaemia (B-ALL Hypo), supratentorial ependymoma (EPD ST), Ewing’s sarcoma (EWS), medulloblastoma group 4 (MB group 4), non-diploid B-cell acute lymphoblastic leukaemia (B-ALL other), infratentorial ependymoma (EPD IT), hepatoblastoma (HB), medulloblastoma SHH subgroup (MB SHH), Wilms tumour (WT), medulloblastoma WNT subgroup (MB WNT), T-cell acute lymphoblastic leukaemia (T-ALL), retinoblastoma (RB), osteosarcoma (OS), medulloblastoma group 3 (MB group 3), high-grade glioma K27wt (HGG Other), adrenocortical carcinoma (ACC), rhabdomyosarcoma (RMS), high-grade glioma K27M (HGG K27M), Burkitt’s lymphoma (BL), atypical teratoid rhabdoid tumour (ATRT), neuroblastoma (NB), embryonal tumours with multilayered rosettes (ETMR). (**b**) Single base substitution signatures in trophoblast clusters. Each column represents one piece of microdissected tissue. (**c**) Bar chart showing the median proportion of substitutions attributable to signature 18. Abbreviations as per (**a**). (**d**) Partial paternal uniparental disomy of 11p detected in two samples, represented by the BAF of SNPs across 11p. Grey denotes SNPs contributed by the father and black by the mother.

### Functional consequences of somatic placental variants

Annotating functional consequences of all somatic variants found in bulk biopsies and trophoblast samples, indicated that most changes were unlikely to have any sequelae (Extended Data Fig.9, Extended Data Table 3). The majority (42/81 unique variants) of copy number changes lay within fragile sites (Extended Data Table 4). Interestingly, two placentas out of 42 harbored copy number neutral loss of heterozygosity (i.e. paternal uniparental disomy) of chromosome 11p15 (**Fig. 4D**). Inactivation of this locus by imprinting or segmental loss underpins a cancer-predisposing overgrowth syndrome, Beckwith-Wiedemann^15^, when it occurs in fetal lineages. It may also be associated with placental disease, as uniparental disomy of 11p15 has been implicated in driving gestational, trophoblast-derived choriocarcinoma^16^.

## DISCUSSION

In this exploration of the somatic genomes of human placentas, we identified genetic bottlenecks at different developmental stages that confined placental tissues genetically. Most prominently, every placental biopsy that we examined represented an independent clonal trophoblast unit, suggesting that mosaicism represents the inherent trophoblast clonal architecture of human placentas. At the earliest stages of embryo development, we identified additional bottlenecks that segregated placental tissues from inner cell mass derived lineages, genetically isolating trophoblast lineages. Together these bottlenecks may represent developmental pathways through which cytogenetically abnormal cells phylogenetically and spatially separate, thereby rendering them detectable by genomic assays utilized in the clinical assessment of chorionic villi. Our findings thus provide plausible, physiological developmental routes through which confined placental trophoblast mosaicism may arise. We suspect that as our understanding of the clonal dynamics of human embryonic lineages grows, we may find additional bottlenecks that account for placental mosaicism affecting mesenchymal lineages also.

The landscape of somatic mutations in placental biopsies was an outlier compared to the other normal human tissues studied to date. In colon^12^, endometrium^13^, esophagus^17^, liver^18^, or skin^19^, clonal fields either represent morphologically discrete, histological units, such as colonic crypts, or clonal expansions associated with oncogenic mutations. In contrast, clonal fields in placental biopsies were “driverless” developmentally acquired expansions that pervaded areas as large as macroscopic biopsies. Furthermore, placental tissues exhibited a comparatively high mutation rate, an unusual predominant mutational signature, and – uniquely for a normal human tissue – frequent copy number changes, reminiscent of some types of human tumours, in particular certain childhood cancers.

There may be several reasons for the distinct somatic features of human placental tissue. Mutagenesis is likely to broadly differ, quantitatively and qualitatively, between fetal and adult life, as has been seen previously^20^, reflecting the unique growth demands and environmental pressures exerted *in utero*. It is also possible that these somatic peculiarities represent the specific challenges that trophoblast lineages undergo during placental growth, such as the approximate threefold rise in the local oxygen tension of blood surrounding the villi between eight and twelve week’s gestation^21^. Finally, it may be conceivable that as a temporary, ultimately redundant organ, some of the mechanisms protecting the somatic genome elsewhere do not operate in placental trophoblasts.

It is possible that genomic placental alterations contribute to the pathogenesis of placental dysfunction, which is a key determinant of the “Great Obstetrical Syndromes”, such as preeclampsia, fetal growth restriction and stillbirth. Previous studies associating confined placental mosaicism with these syndromes have yielded conflicting results^3,10,22–24^. Our studies may explain these discrepancies, as the genomic alterations we observed were not uniformly distributed across multiple biopsies from the same placenta. Larger scale systematic studies of the genomic architecture of the human placenta in health and disease might establish the role of placental genomic aberrations in driving placenta-related complications of human pregnancy.

## METHODS

### Ethics statement

All the samples were obtained from the Pregnancy Outcome Prediction (POP) study, a prospective cohort study of nulliparous women attending the Rosie Hospital, Cambridge (UK) for their dating ultrasound scan between January 14, 2008, and July 31, 2012. The study has been previously described in detail^25,26^. Ethical approval for this study was given by the Cambridgeshire 2 Research Ethics Committee (reference number 07/H0308/163) and all participants provided written informed consent.

### Bulk DNA sequencing

DNA was extracted from maternal blood, umbilical cord, and fresh frozen placental biopsies. Short insert (500bp) genomic libraries were constructed, flow cells prepared and 150 base pair paired-end sequencing clusters generated on the Illumina HiSeq X or NovaSeq platform according to Illumina no-PCR library protocols. An overview of samples and sequencing variables, including the average sequence coverage, is shown in Extended Data Table 1.

### Laser capture microdissection and low-input DNA sequencing

Tissues were prepared for microdissection and libraries were constructed as described previously^12,13^ and subsequently submitted for whole-genome sequencing on the Illumina HiSeq X or NovaSeq platform.

### DNA sequence alignment

All DNA sequences were aligned to the GRCh37d5 reference genome by the Burrows-Wheeler algorithm (BWA-MEM)^27^.

### Detection of somatic variants

We called all classes of somatic mutations: substitutions (CaVEMan algorithm^28^, see below), indels (Pindel algorithm^29^), copy number variation (ASCAT^30^ and Battenberg^12,13^ algorithms), and rearrangements (BRASS algorithm^12,13^). Besides ASCAT and Battenberg, sub-chromosomal copy number variants can also be picked up via the breakpoints as predicted by BRASS, providing three independent methods to call copy number variants. The umbilical cord sample functioned as a matched normal sample in variant calling.

Rearrangements were validated by local assembly (as implemented in the BRASS algorithm). To generate a high confidence, final list of structural variants, only rearrangements whose breakpoints were greater than 1,000 base pairs apart, absent in the germline and associated with a copy number change were included in our analysis (see Extended Data Table 4). All calls were visually inspected in the genome browser Jbrowse^31^.

### Unmatched substitution calling

Substitutions were called by applying the CaVEMan^28^ algorithm in an unmatched analysis of each sample against an *in silico* human reference genome. Beyond the inbuilt post-processing filter of the algorithm, we removed variants affected mapping artefacts associated with BWA-MEM by setting the median alignment score of reads supporting a mutation as greater than or equal to 140 (ASMD>=140) and requiring that fewer than half of the reads were clipped (CLPM=0). We then recounted across samples belonging to the same patient the variant allele frequency of all substitutions with a cut-off for base quality (=25) and read mapping quality (=30). Variants were also filtered out if they were called in a region of consistently low or high depth across all samples from one patient.

To filter out germline variants, we fitted a binomial distribution to the combined read counts of all normal samples from one patient per SNV site, with the total depth as the number of trials, and the total number of reads supporting the variant as number of successes. Germline and somatic variants were differentiated based on a one-sided exact binomial test. For this test, the null hypothesis is that the number of reads supporting the variants across copy number normal samples is drawn from a binomial distribution with p=0.5 (p=0.95 for copy number equal to one), and the alternative hypothesis drawn from a distribution with p<0.5 (or p<0.95). Resulting p-values were corrected for multiple testing with the Benjamini-Hochberg method and a cut-off was set at q < 10^−5^ to minimize false positives as on average, roughly 40,000 variants were subjected to this statistical test. Variants for which the null hypothesis could be rejected were classified as somatic, otherwise as germline.

Further, remaining artefacts were filtered out by fitting a beta-binomial distribution to the variant counts and total depth for all variants across all samples from one patient. From this set of observations, we quantified the overdispersion parameter (rho). Any variant with an estimated rho smaller than 0.1 was filtered out, as used previously^32,33^.

Following visual inspection of a subset of these putative variants using Jbrowse^31^, a small number of substitutions called within the placental biopsies were found to falsely pass at sites of germline indels. To remedy this, substitutions called at the site of an indel were removed.

### Phylogeny reconstruction

Phylogenies of microdissected trophoblast clusters were generated from the filtered substitutions using a maximum parsimony algorithm, MPBoot^34^. Substitutions were mapped onto tree branches using a maximum likelihood approach.

### Unmatched indel calling

A similar approach was taken for indel filtering. Variants in each sample were called against the *in silico* human reference genome using Pindel^29^. Those that passed and possessed a minimum quality score threshold (>=300) were subject to the same genotyping and fitting of binomial and beta-binomial distributions described above and only variants supported by at least five mutant reads were retained.

Some samples with higher coverage (>50X) retained an inflated number of low VAF indel calls following this filtering approach. Further investigation revealed that most of these excess calls to occur at sites Pindel frequently rejects in other unrelated samples sequenced using the same sequencing platforms, suggesting that they were artefactual in nature. As these samples accounted for the majority of low VAF indels called in the biopsies, indels with a VAF <0.1 in these bulk samples were removed. Again, a subset of called indels were reviewed in Jbrowse^31^ to check the veracity of the pipeline detailed here.

### Exclusion of maternal contamination

To exclude the possibility of any remaining maternal DNA in the placenta to skew results on mutation burden and clonality, we used maternal SNPs to quantify contamination. For each pregnancy, we randomly picked 5,000 rare germline variants (i.e. left in by the common SNP filter in CaVEMan) found in mother but not in umbilical cord. All these variants passed other CaVEMan flags, did not fall in regions of low depth (on average, below 35), and were present at a VAF greater than 0.35 in mother. Their VAFs in all individual placental samples, microdissections and biopsies, is displayed in Extended Data Fig.1. No sample had a level of support for maternal SNPs that exceeded the expectations for sequencing noise (0.1%), excluding maternal contamination as a plausible origin for any observations made here.

### Sensitivity correction of mutation burden

To compensate for the effects of sequencing coverage and low clonality on the final mutation burden per sample, we estimated the sensitivity of variant calling. For each sample, we generated an *in silico* coverage distribution by drawing 100,000 times from a Poisson distribution with the observed median coverage of the sample as its parameter. For each coverage simulation, we calculated the probability of observing at least four mutant reads for SNVs or five for indels (the minimum depth requirement for our CaVEMan and Pindel calls respectively) with the underlying binomial probability given by the observed median VAF of the sample. The average of all these probabilities then represents the sensitivity of variant calling. Final mutation burdens were then obtained by dividing the observed number of mutations by the estimated sensitivity.

### Mutational signature extraction and fitting

To identify possibly undiscovered mutational signatures in human placenta, we ran the hierarchical Dirichlet process (HDP) (https://github.com/nicolaroberts/hdp) on the 96 trinucleotide counts of all microdissected samples, divided into individual branches. To avoid overfitting, branches with fewer than 50 mutations were not included in the signature extraction. HDP was run with individual patients as the hierarchy, in twenty independent chains, for 40,000 iterations, with a burn-in of 20,000.

Besides the usual flat noise signature (Component 0) that is usually extracted, only one other signature emerged (Component 1) from the signature extraction. Deconvolution of that signature revealed it could be fully explained by a combination of reference signatures SBS1, SBS5, and SBS18 (Extended Data Fig.9), all of which have been previously reported in normal tissues.

Because of the lack of novel signatures in this data set, the remainder of mutational signature analysis was performed by fitting this set of three signatures to trinucleotide counts using the R package deconstructSigs (v1.8.0)^35^.

### Genetic proximity scores

To measure the genetic proximity between any two trophoblast clusters from the same biopsy, we used the following equation:

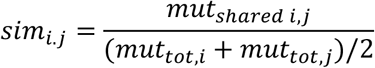

Or simply, the fraction of shared mutations between samples *i* and *j* divided by their average total mutation burden. The resulting number reflects how much of *in utero* development was shared between these samples.

However, control data of normal human colon^12^ and endometrium^13^ were obtained from adults and their phylogenetic histories will reflect postnatal tissue dynamics as well. To obtain a proxy for the sharedness due to development *in utero*, we only considered a pair of samples *i* and *j,* if they did not split at a mutational time inconsistent with early development. We set this threshold for both colon and endometrium at 100 mutations, a very rough estimate of the maximum burden at birth in these tissues given preliminary studies. Consequently, instead of dividing the number of early shared mutations by the average burden, for adult tissues, these were divided by 100.

### Embryonic mutations

To discover early mutations in the umbilical cord samples, we included these in the unmatched variant calling as described above, either with all bulk placenta samples or microdissections. In the case of the latter, the umbilical cord samples were not included in phylogeny reconstruction due to their polyclonality, but aggregating it with microbiopsy data allows for effective removal of germline variants due to the high cumulative depth of coverage.

All embryonic variants were visually inspected in Jbrowse^30^ to exclude any possible remaining sequencing or mapping artefacts.

For the five phylogenies of trophoblast clusters, the contribution of branches to the umbilical cord was measured by the VAF of mutations on these branches. In PD42138 and PD42142, where no variants were shared between the trophoblast phylogeny and the umbilical cord, the earliest mutations were found exclusively in the umbilical cord sample and the mutations with the highest VAF were taken to delineate the major clone, as done for sets of bulk biopsies. In both cases, the VAFs of the earliest mutations reflected a clonal origin for umbilical cord.

For bulk placenta samples and umbilical cord, the asymmetric contribution of the zygote was calculated by converting the highest VAF found in umbilical cord to a contribution (effectively multiplying by two). The alternative lineage was identified using the pigeonhole principle^13^, i.e. when clustering of the VAFs across placenta and umbilical cord prohibited this lineage from being a sub-clone of the previously identified major clone. In about half of cases (17/37), this yielded an asymmetry in umbilical cord with major and minor lineage also fully accounting for the placental bulk samples (see Extended Data Fig.4). For one case (PD45595), we could not identify any non-artefactual early embryonic mutations in the umbilical cord. This patient is hence omitted from the subsequent analysis concerning the early asymmetries.

In 11 out of 37 cases (Extended Data Fig. 5), one or more of the placental lineages could not be fully explained by the umbilical cord lineages, although the latter exhibited the expected asymmetry. This was established by calculating the 95% confidence intervals around the expected binomial probabilities of both major and minor lineages. If the sum of the higher extremes was less than 0.5 (the expected value to fully account for this lineage), the placental biopsy was not fully explainable by the umbilical cord lineages.

In the remaining 9 out of 37 cases (Extended Data Fig.6), the umbilical cord showed clonal origins (a major lineage with a VAF around 0.5), which we found to be paired with segregated placental lineages in all cases.

### Genotyping germline SNPs on chromosome 10

PD45581c, a bulk placenta biopsy, exhibited trisomy of chromosome 10, which was absent from PD45581e (placenta) and PD45581f (umbilical cord). This could be the result of either a somatic duplication of chromosome 10 or a trisomy present in the fertilised egg that was post-zygotically reverted to a disomy. These two scenarios can be distinguished from one another by the number of distinct chromosomal alleles: three different chromosomes for a trisomic rescue, two for a somatic duplication. To test this, all SNPs on chromosome 10 reported by the 1000 Genomes project were genotyped across the three samples from the pregnancy, as well as the mother.

### Coding substitution rate of trophoblast clusters against paediatric cancers

A recent, large scale, pan-paediatric cancer project provided the data necessary to contrast against the high mutation rate we observe in the trophoblast^14^. Here, the burden analysis focused on ‘coding mutations’, taken to mean all SNVs and indels that lie within exonic regions. This was adjusted for the callability and expressed per megabase.

To generate comparable results from our data, we used mosdepth (https://github.com/brentp/mosdepth) to estimate the callable length of the autosomal exonic regions. This meant excluding all regions blacklisted during variant calling, such as those with more mappability, and those with insufficient sequencing depth to call substitutions (<4X). Our substitution burden estimates were then divided by our percentage estimate of the autosomal exonic regions covered. To compare the rate of mutagenesis rather than gross burden, this figure was then divided by the age (in years) plus 0.75. This would adjust for gestation and any substitutions gained in the tumour precursor whilst still in utero.

### Calculating the burden of SBS18 compared to paediatric malignancies

Using only the tumours that had undergone whole genome sequencing and SBS signature extraction in the paper listed above, we simply expressed the SBS18 mutations as a proportion of all SNVs and ranked the median value returned per tumour against the trophoblast clusters.

### Chromosome 11p phased B-allele frequency plotting

ASCAT and Battenberg identified two samples, PD45557e_lo0003 and PD42154b3, as having uniparental disomy of part of chromosome 11p. To phase this to a given parent, all SNPs identified by the 1000 Genomes project on chromosome 11p were genotyped for the affected sample, the matched umbilical cord and the maternal blood sample. The SNPs that were homozygous in the mother but heterozygous in the umbilical cord could then be used to phase the loss of heterozygosity in the placental sample as the remaining allele must belong to the father.

## DATA AVAILABILITY

DNA sequencing data are deposited in the European Genome-Phenome Archive (EGA) with accession code EGAD0000100637.

## CODE AVAILABILITY

Bespoke R scripts used for analysis and visualization in this study are available online from GitHub (https://github.com/TimCoorens/Placenta).

## ACKNOWLEDGEMENTS

We thank Professor Sir Michael Stratton and Dr Iñigo Martincorena for discussions.

## CONTRIBUTIONS

S.B. designed the experiment. T.H.H.C. performed phylogenetic analyses. T.H.H.C. and T.R.W.O. analysed somatic mutations. T.R.W.O. performed microdissections. R.S., U.S., E.C., R.V.-T., M.H., M.D.Y., and R.R. contributed to experiments or analyses. N.S. provided pathological expertise. P.J.C. contributed to discussions. S.B., T.H.H.C. and T.R.W.O. wrote the manuscript, aided by D.S.C.J. and G.S. D.S.C.J., G.S., and S.B. co-directed this study.

## COMPETING INTERESTS

No competing interests are declared by the authors of this study.

**Extended Data Figure 1.**
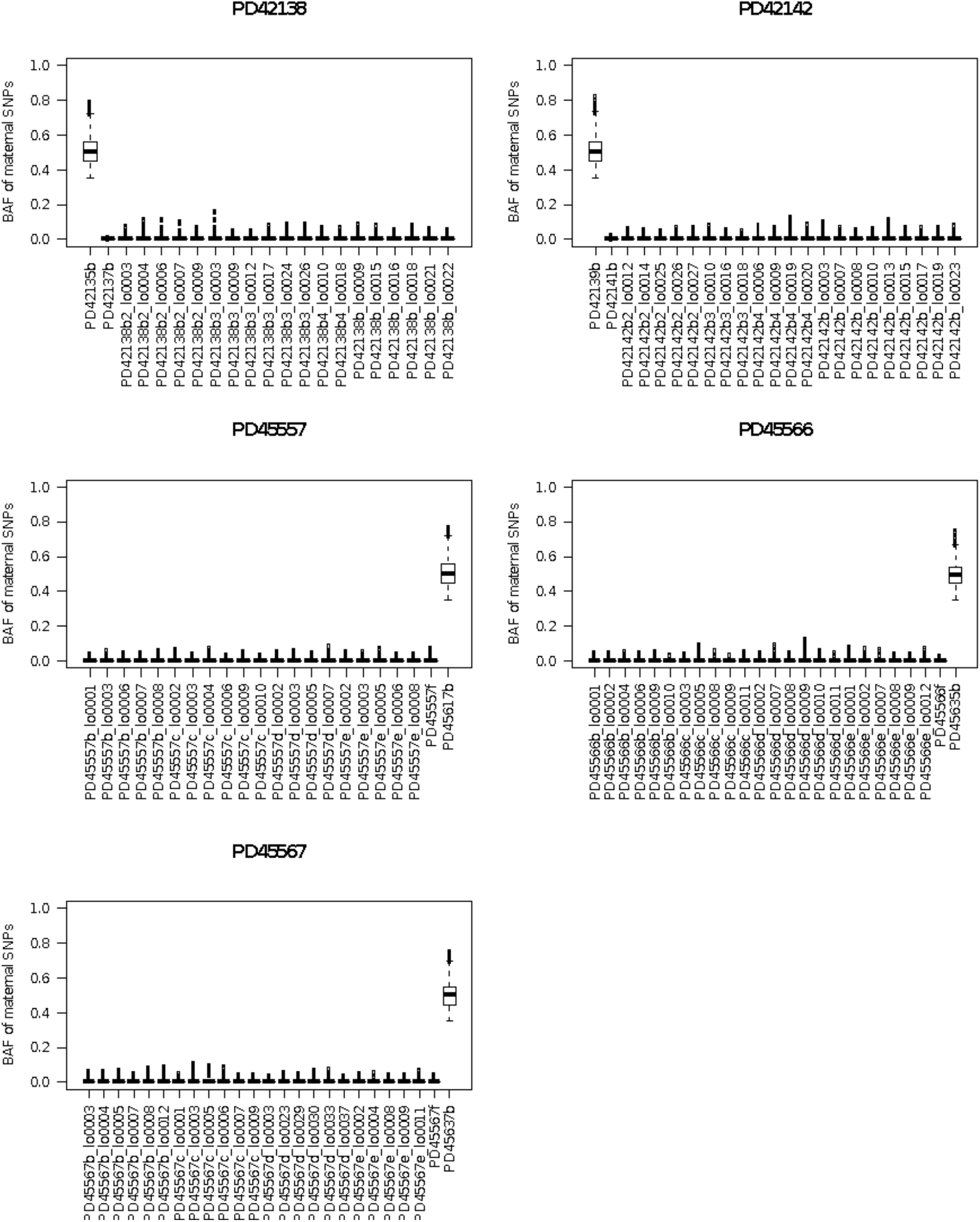
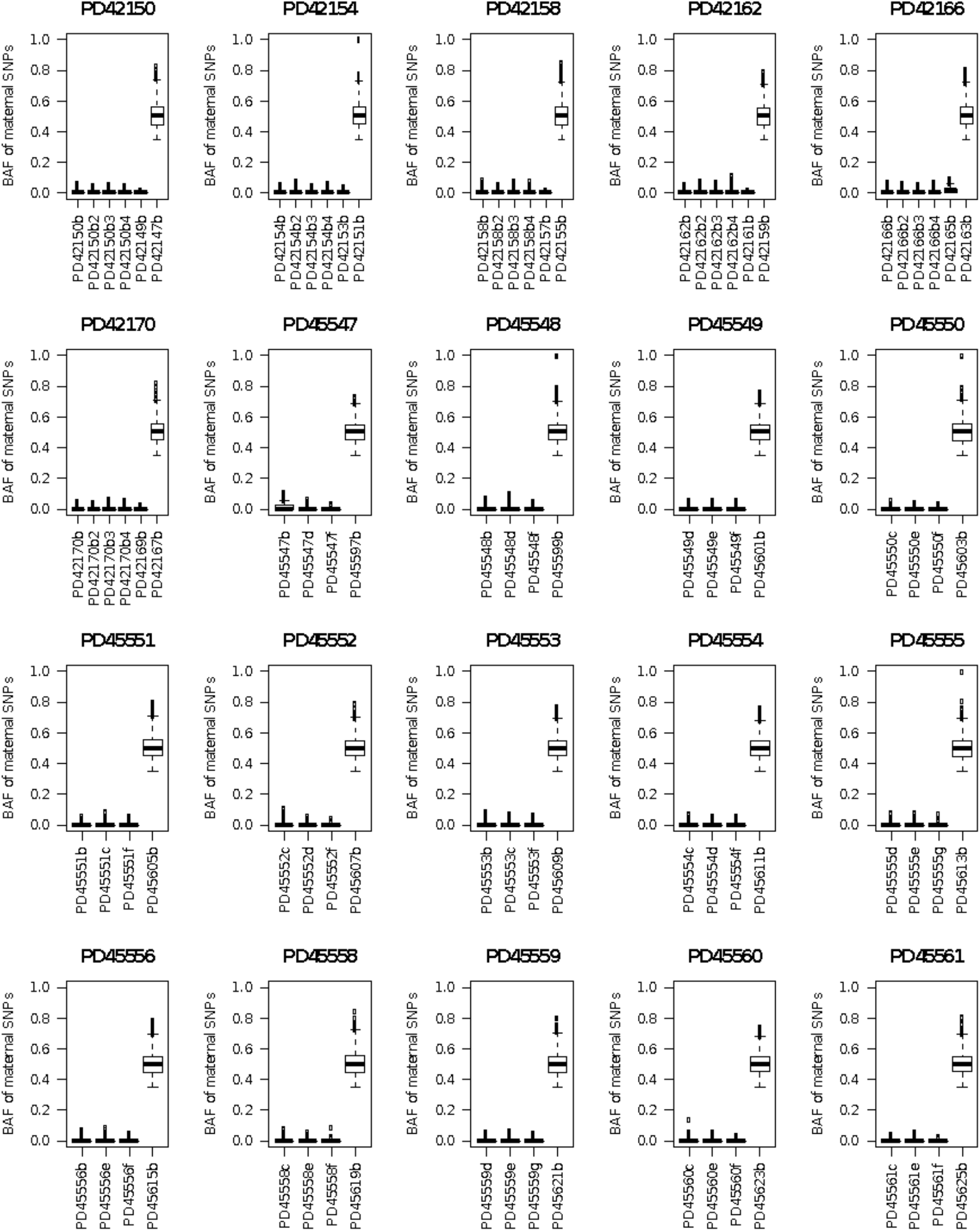
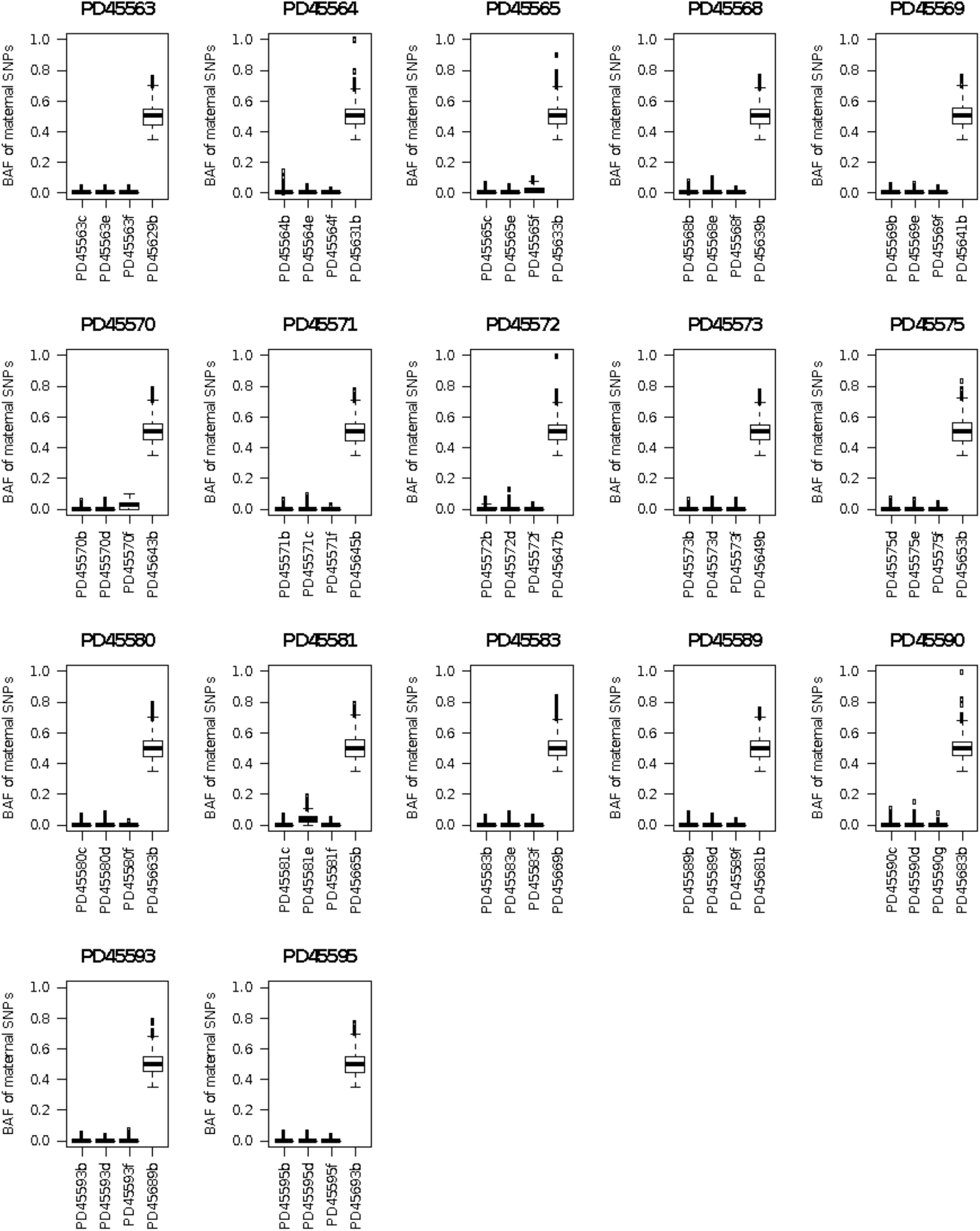
Exclusion of maternal contamination. Boxplots of B-allele frequency (BAF) of rare SNPs called in mother, but absent from umbilical cord, as an indicator of possible maternal contamination across placental samples. The maternal blood sample is placed in each plot (furthest right) as a control.

**Extended Data Figure 2.**
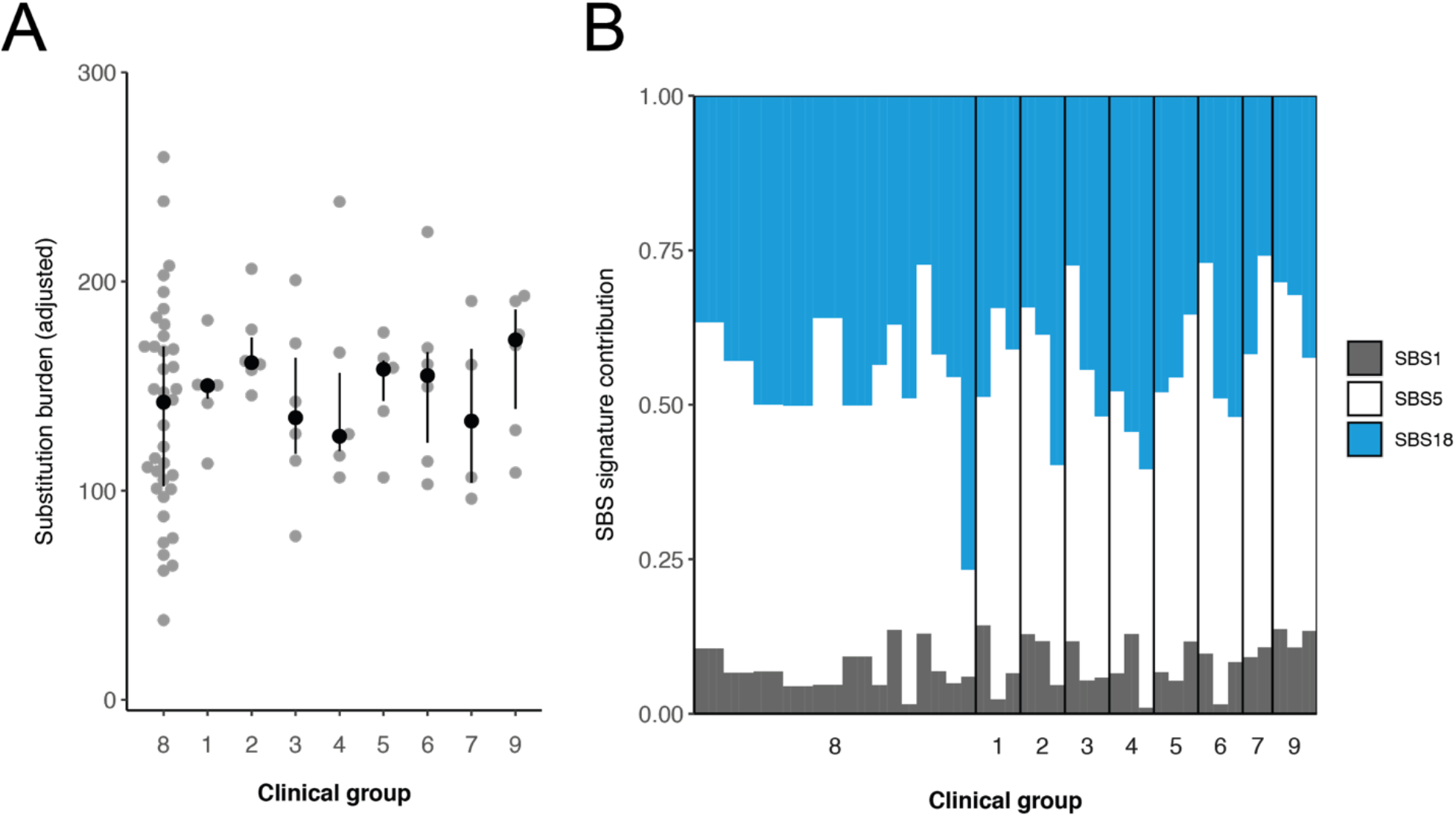
Differences in substitutions between clinical groups. Analysis per clinical group of the absolute substitution burden of each placental biopsy (A) and their associated mutational signatures (B). The difference in substitution burden between the clinical groups is not significant (Kruskal-Wallis rank sum test, p=0.7438). Each point and bar represent a single placental biopsy. Clinical groups are defined in table S1.

**Extended Data Figure 3.**
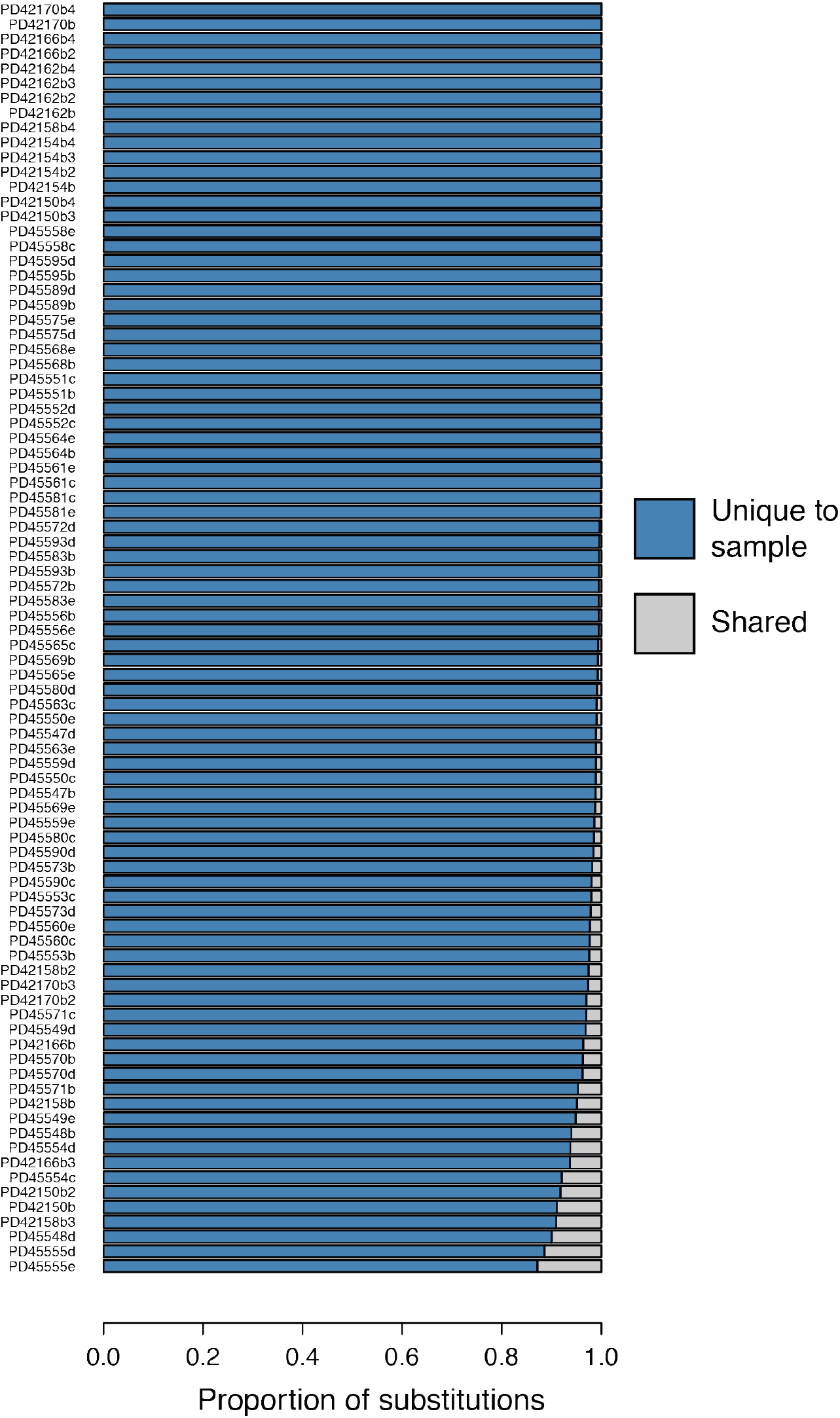
Unique variants in placental biopsies. Proportion of variants that are unique to each placental biopsy (blue), so absent from matched umbilical cord as well as any other placental biopsy sampled from the same patient.

**Extended Data Figure 4.**
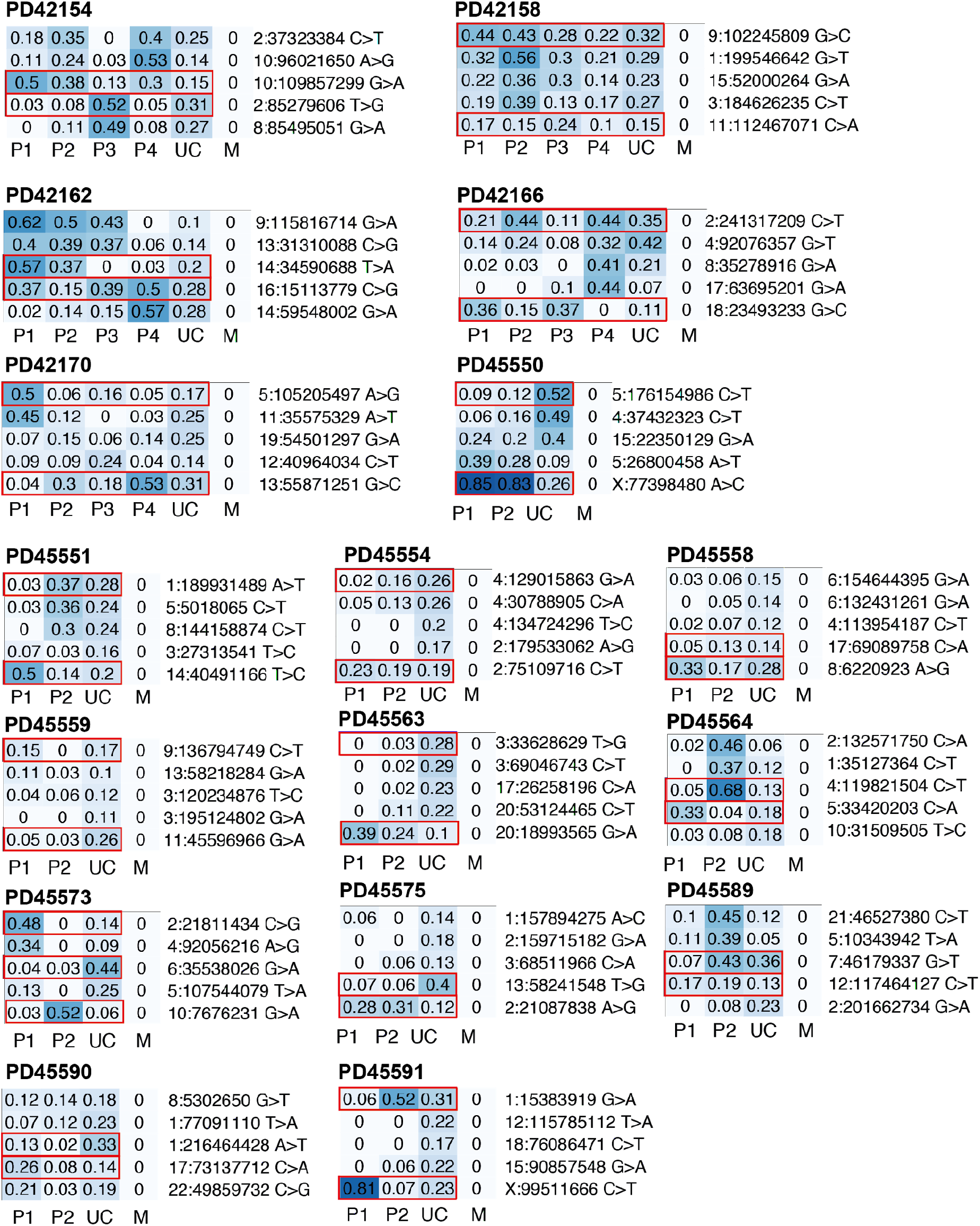
Asymmetry across trophectoderm and umbilical cord. Heatmaps of VAFs of early embryonic mutations with the two earliest lineages contributing both to placenta and umbilical cord. Putative earliest mutations highlighted in red. (P=placenta, UC=umbilical cord, M=maternal)

**Extended Data Figure 5.**
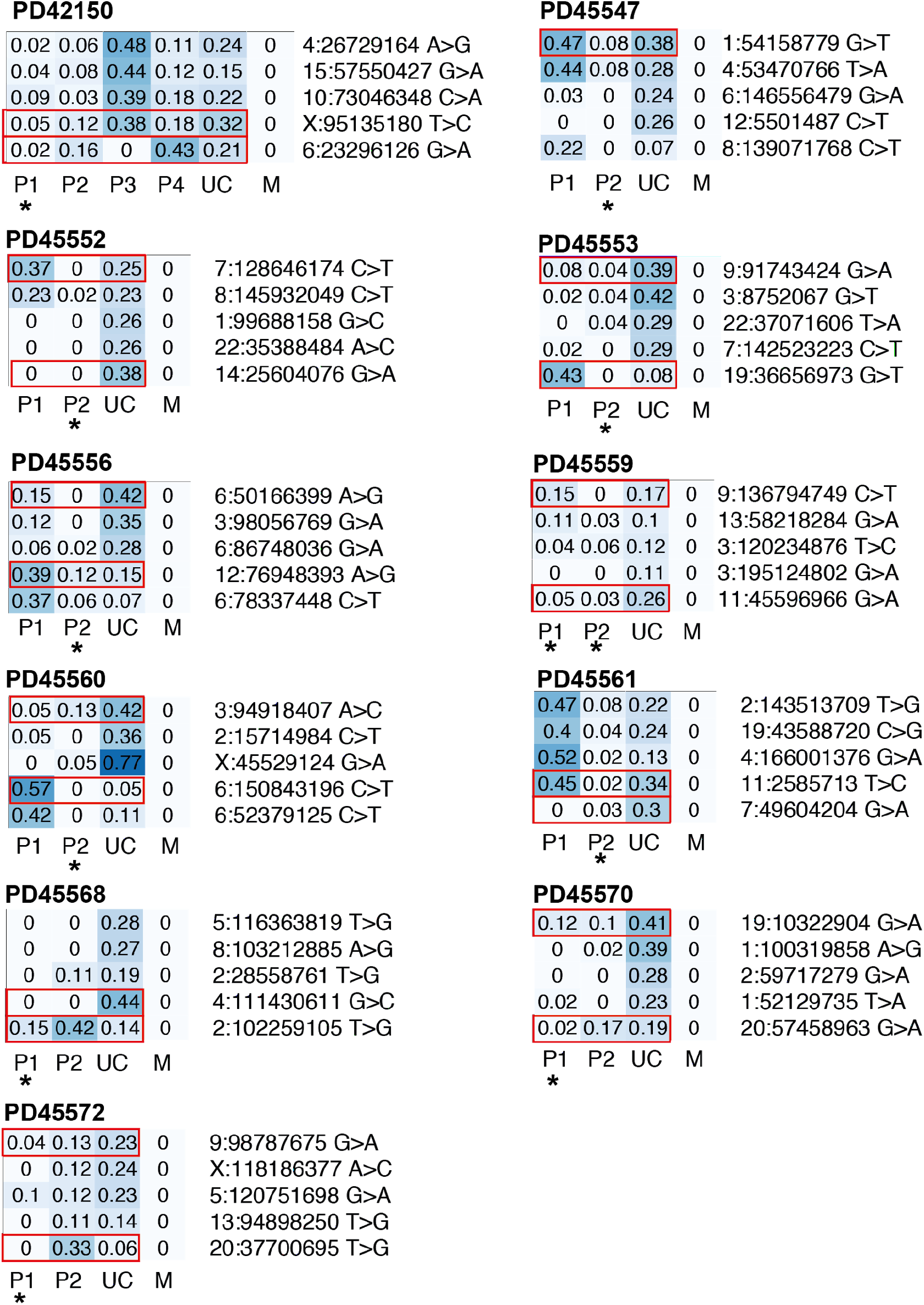
Unexplained placental lineages. Heatmaps of VAFs of early embryonic mutations with the two earliest lineages contributing umbilical cord. Putative earliest mutations highlighted in red. Asterisk indicates placental lineage is not fully explained by umbilical cord (see Methods). (P=placenta, UC=umbilical cord, M=maternal)

**Extended Data Figure 6.**
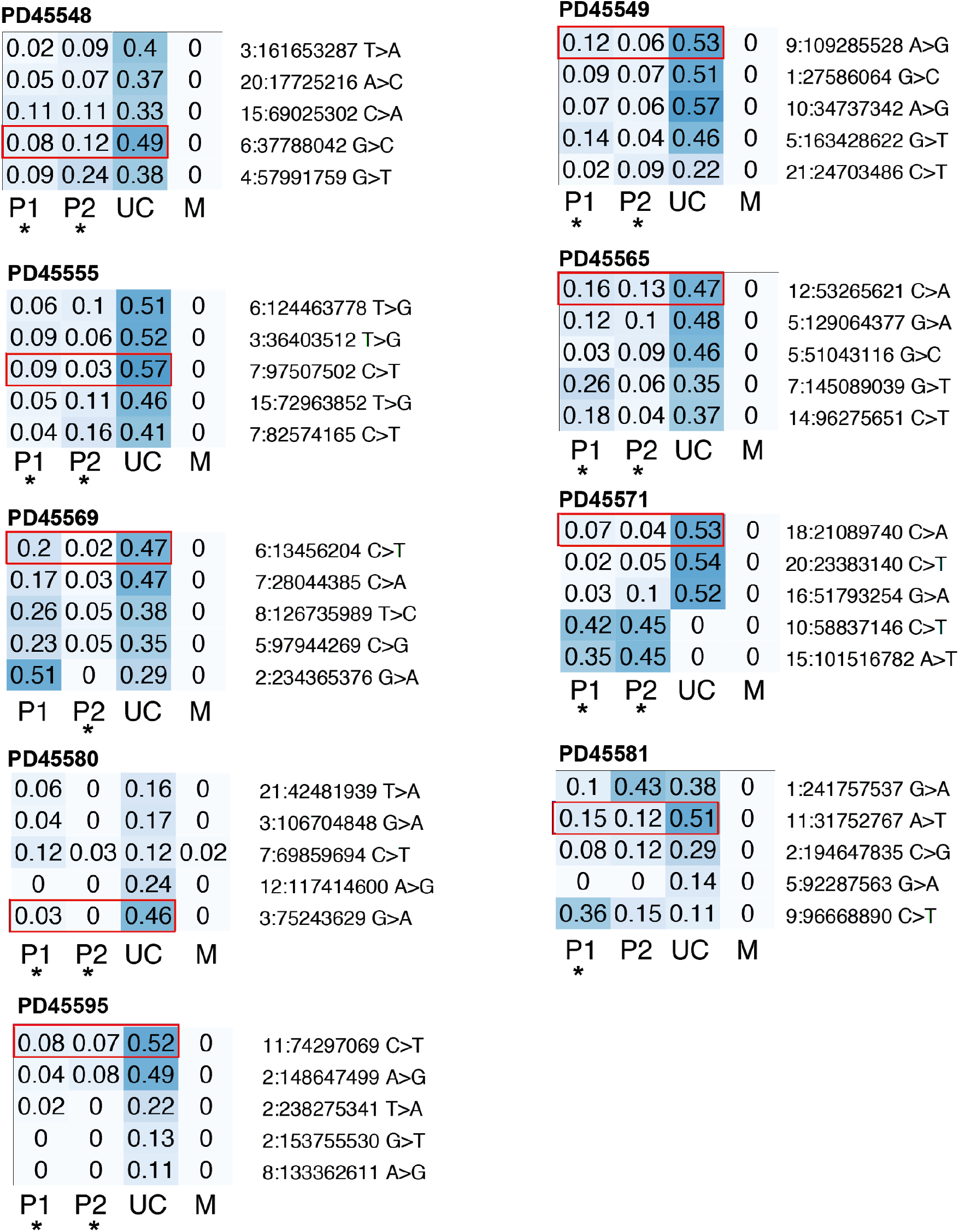
Full segregation of placental and umbilical cord lineages. Heatmaps of VAFs of early embryonic mutations with the umbilical cord being derived from one clonal lineage. In all cases, one or more placental lineages do not share any genetic ancestry with umbilical cord and are largely unexplained, as indicated by an asterisk (see Methods). (P=placenta, UC=umbilical cord, M=maternal)

**Extended Data Figure 7.**
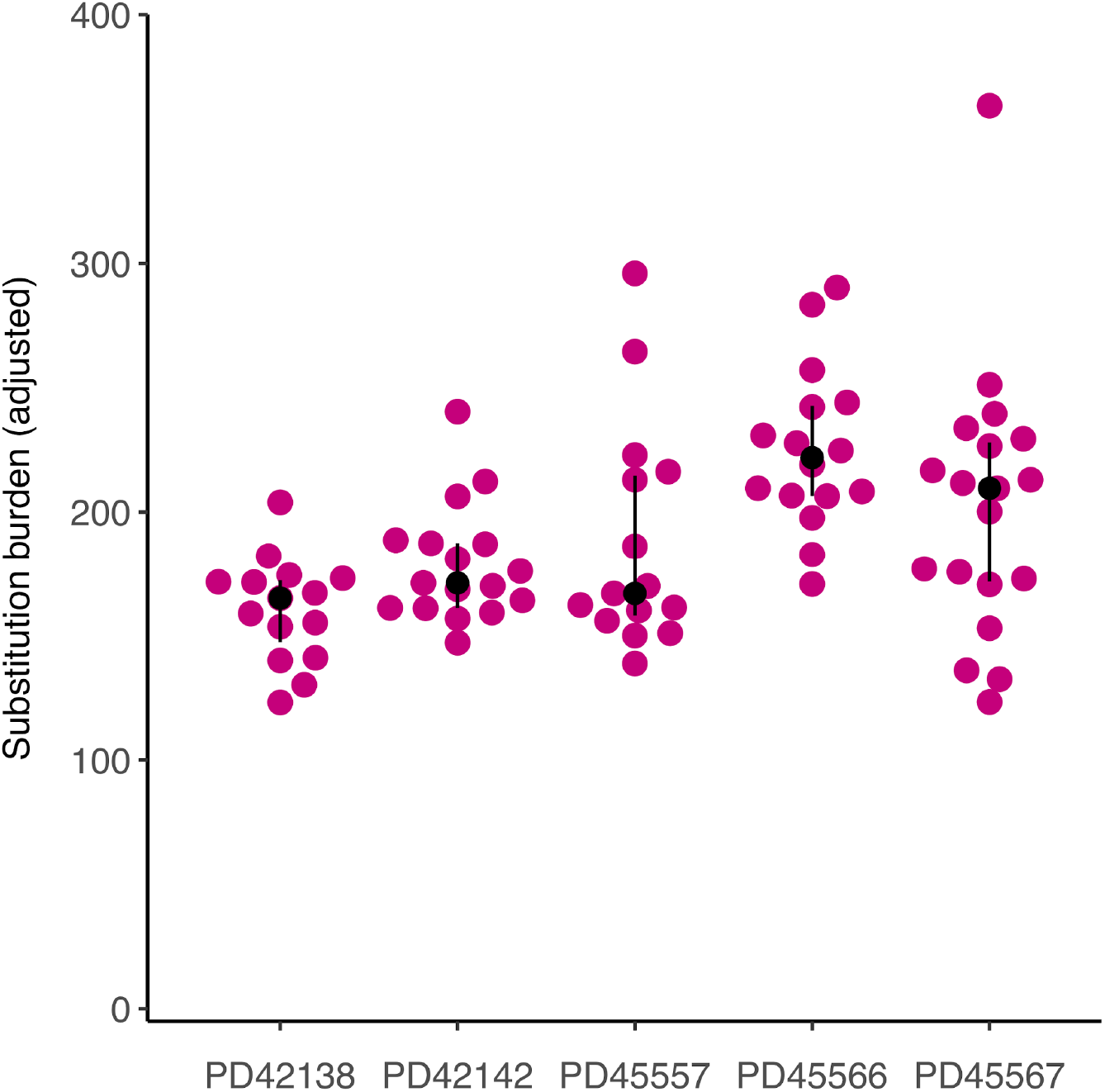
Substitution burden per individual trophoblast cluster. Adjusted for coverage and median variant allele frequency.

**Extended Data Figure 8.**
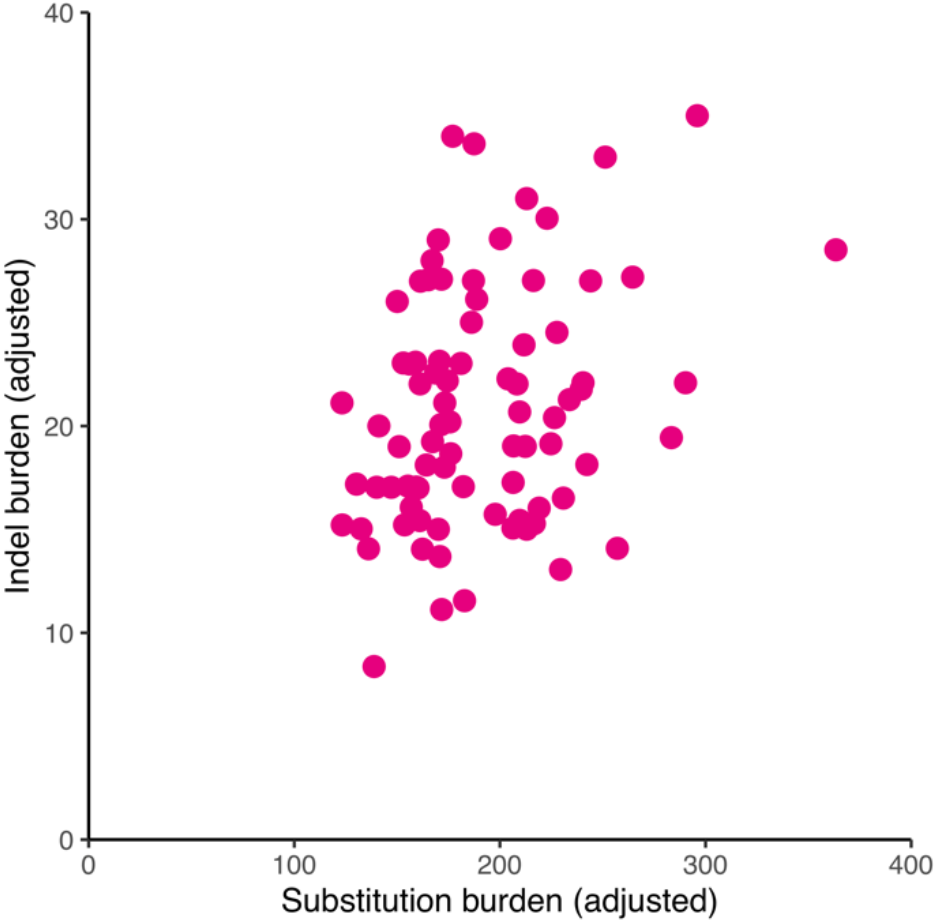
Indels versus substitutions. Indel burden versus substitution burden per trophoblast cluster. Both are corrected for median VAF and coverage.

**Extended Data Figure 9.**
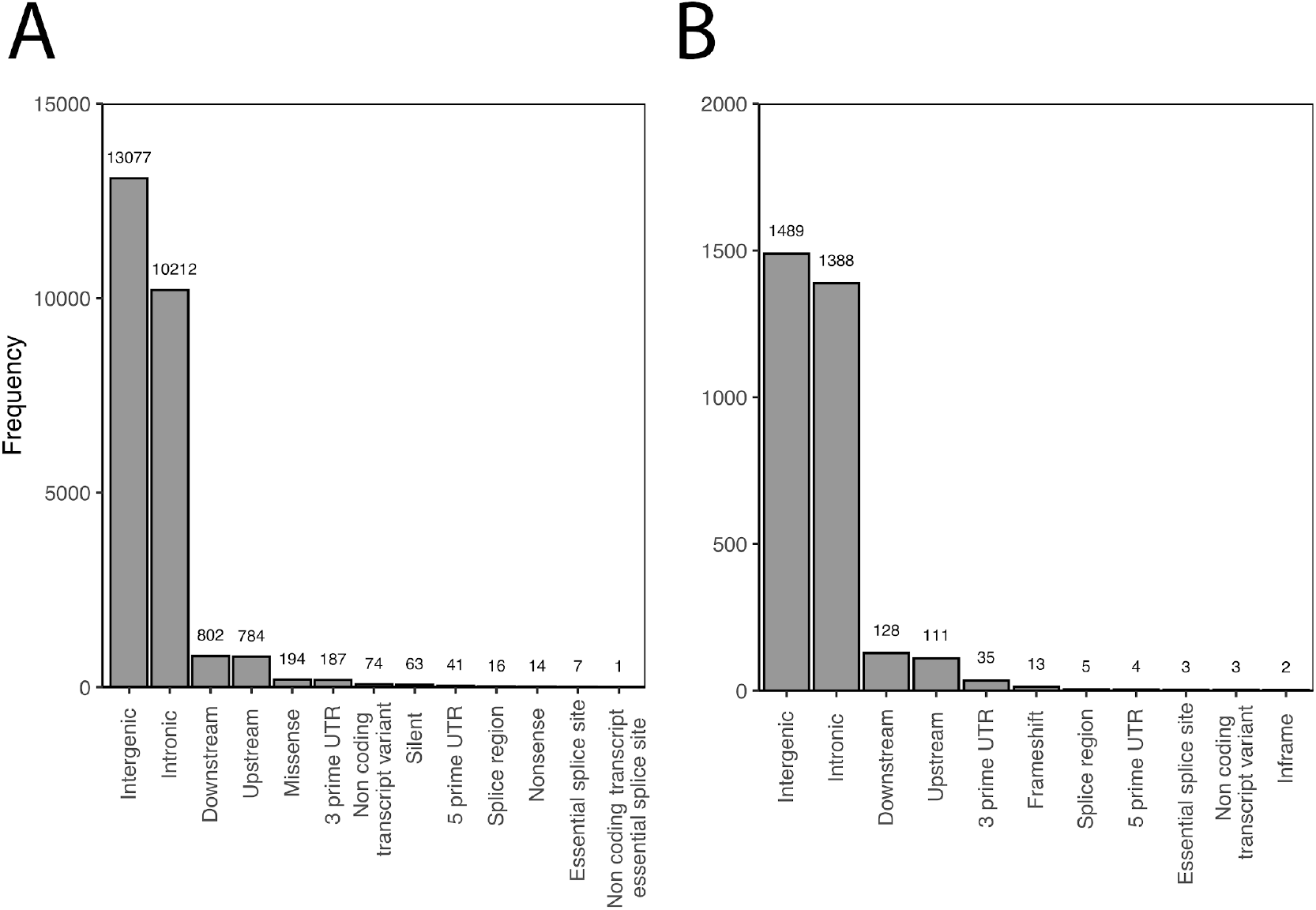
Impacts of mutations. Overview of functional consequences of unique SNVs (**A**) and indels (**B**) seen in the placental biopsies and trophoblast clusters.

**Extended Data Figure 10.**
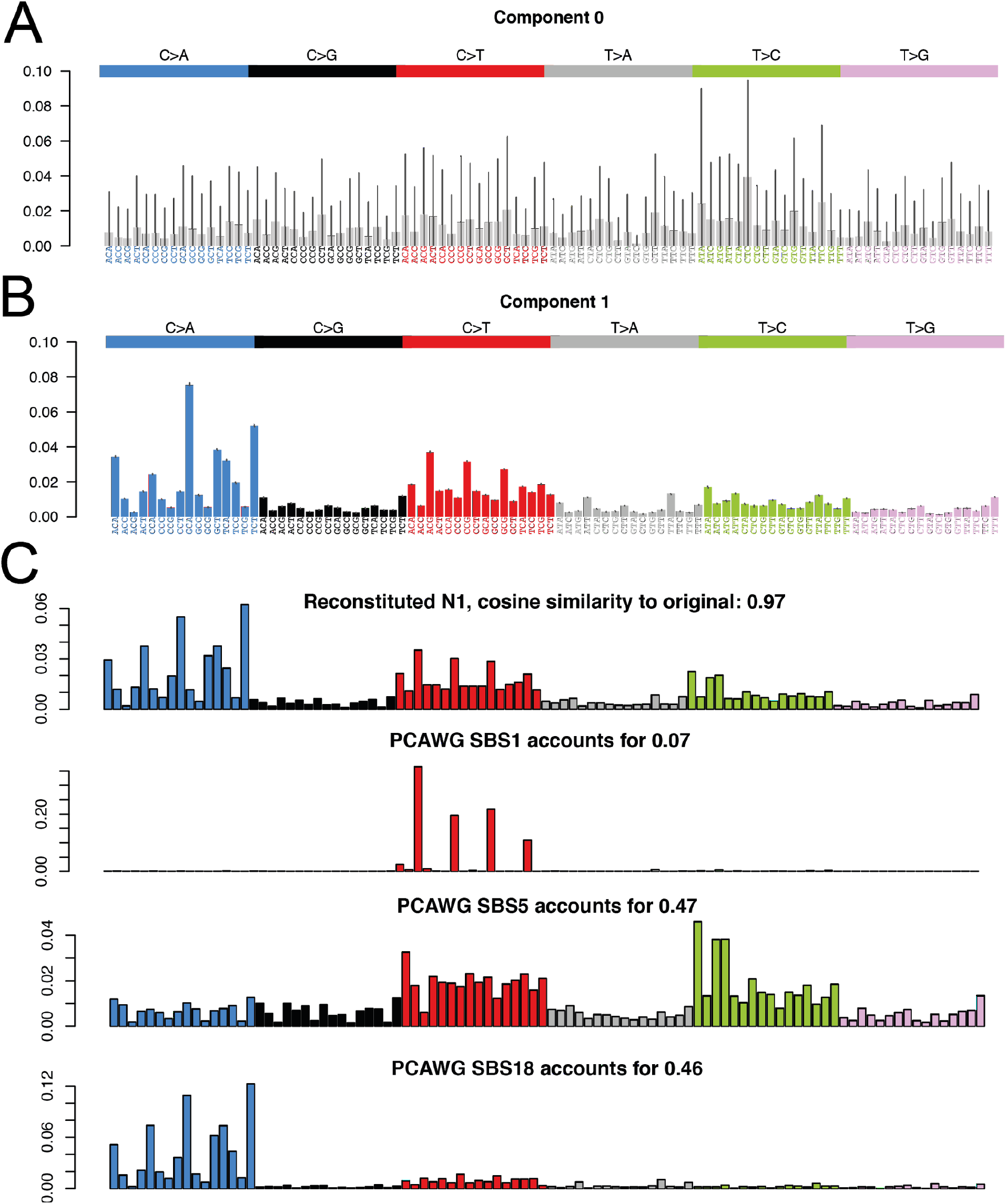
Signatures extraction and deconvolution. Signature extraction by HDP yielded a noise component (**a**) and one genuine mutational signature (**b**), which could be convoluted and reconstructed using three reference mutational signatures: SBS1, SBS5 and SBS18 (**c**).

